# Dual Targeting for Enhanced Tumor Immunity: Conditionally Active CD28xVISTA Bispecific Antibodies Promote Myeloid-Driven T-Cell Activation

**DOI:** 10.1101/2025.04.07.647657

**Authors:** Thomas Thisted, F. Donelson Smith, Zhi-Gang Jiang, Zuzana Biesova, Adejumoke Onumajuru, Yuliya Kleschenko, Kanam Malhotra, Vikas Saxena, Arnab Mukherjee, Edward H. van der Horst

**Author notes:** Corresponding author: Edward H. van der Horst, Sensei Biotherapeutics Inc., 1405 Research Blvd., Suite 125, Rockville MD 20850. **Authors’ Disclosures:** T.T., F.D.S., Z-G.J., Z.B., A.O., Y.K., K.M., V.S., A.M, and E.H.vdH are current or former employees of Sensei Biotherapeutics, Inc. The authors declare no other conflicts of interest.

## Abstract

Reinvigoration of tumor-reactive T-cells using co-stimulatory bispecific antibodies (bsAbs) targeting CD28 or CD137 is emerging as a promising therapeutic strategy. Conditional, tumor-specific recruitment can offer a necessary layer of control and specificity. We developed pH-selective CD28xVISTA bsAbs to act specifically within the acidic tumor microenvironment (TME), aiming for enhanced T-cell-mediated cancer cell killing while minimizing systemic T-cell activation and Cytokine Release Syndrome (CRS) risk. CD28 agonism by CD28xVISTA bsAbs relies on pH-selective engagement of VISTA, a protein robustly expressed on myeloid cells highly prevalent in most solid tumors. This modality avoids engagement of tumor-associated antigens (TAAs) with the potential to provide highly tumor specific activity with minimal on-target/off-tumor side effects.

We report the identification of a lead candidate with pH-dependent simultaneous engagement of both targets, and VISTA-dependent CD28 signaling in a reporter cell line. CD28xVISTA avidly bound VISTA-positive cells, and co-stimulation was shown *in vitro* by its ability to activate and expand T-cells and enhance T-cell mediated cancer cell killing in co-cultures of human PBMCs and cancer cells in the presence of a TAA-targeted anti-CD3 T-cell engager. Interestingly, our findings support both signaling in *cis* (between T-cell and cell displaying peptide-MHC complex) and in *trans* with stimulation occurring through CD28 clustering outside of the immune synapse. Our lead candidate displayed efficient tumor growth inhibition of human VISTA-expressing MC38 cells in a humanized CD28 syngeneic mouse model in combination with PD-1 blockade. Importantly, our CD28xVISTA bsAb showed no signs of superagonistic properties in several *in vitro* assays geared towards revealing induction of CRS. Our data supports clinical development in combination with anti-PD-1 or any TAA-targeted anti-CD3 T-cell engagers developed for solid tumors.

## INTRODUCTION

T-cell activation is initiated when the T-cell receptor/CD3 complex (TCR/CD3) binds to immunogenic peptides in the context of the major histocompatibility complex (MHC) expressed by professional antigen-presenting cells (APCs), virally infected cells, or tumor cells. This initial TCR/peptide MHC (pMHC) signal for T-cell activation has been termed “Signal 1”. TCR/CD3 complexes cluster together at the interface between T-cells and their target cells called the “immune synapse”. By itself, Signal 1 is not sufficient to achieve full T-cell stimulation, but additional co-stimulatory signals (“Signal 2”) and further cytokine stimulation is required for maximal T-cell activation. Co-stimulatory receptors such as CD28 or CD137 (4-1BB) also accumulate and interact with their respective ligands at the immune synapse. The profound role of the interaction between CD28 and its cognate ligands CD80 and CD86 (a.k.a. B7.1 & B7.2) was first highlighted by the induced expression of B7 on mouse melanoma cells leading to their complete rejection by CD8^+^ T-cells *in vivo* (1). Tumor cells do not typically express CD80 & CD86 (these are expressed primarily on APC’s), and this lack of co-stimulation may compromise the activity of CD3 engagers or anti-PD1 therapies in the clinic. However, using bsAbs with one arm engaging a tumor-associated antigen (TAA) and the other binding to CD28 on T-cells can artificially bridge TAAs on tumor cells to CD28 receptors on T-cells in the context of the synapse (i.e. in *cis*), causing CD28 clustering and activation. Preclinical efficacy for co-stimulatory bsAbs was first shown for CD28 (2,3), and T-cell co-stimulation through 4-1BB or CD28 is currently being explored in the clinic (4,5).

In contrast to hematological cancers, highly specific TAA’s for solid tumors are rare (6). Low level TAA expression on normal tissue can lead to on-target/off-tumor toxicities. All disclosed bi-and multi-specific CD28-targeted mAbs (currently over 20 in pre-clinical or clinical development (5)), rely on engaging with at least one TAA. Development of immunostimulatory antibodies through receptor agonism without associated toxicity is particularly prudent in the context of CD28, where superagonism (i.e. full T-cell activation in the absence of TCR ligation) can lead to severe life-threatening toxicity in the form of cytokine release syndrome (CRS; (7)). Therefore, an alternative mode of action that circumvents TAA’s but provides a high degree of tumor specificity could be highly advantageous.

VISTA (V-domain Ig-containing suppressor of T-cell activation) is highly expressed on tumor abundant myeloid cells, including antigen-presenting cells (APCs; e.g. macrophages and dendritic cells (DCs)). It plays a role in modulating the immune response by functioning as a negative regulator of the immune system. However, unlike other immune checkpoints (e.g. PD-1 or CTLA-4), VISTA is only active in the low pH environment (∼pH 6) found inside many tumors. This is brought about by pH-selective engagement of the inhibitory receptor P-selectin glycoprotein ligand-1 (PSGL-1) on T-cells (8,9).

Here we exploit the abundant tumor infiltration by VISTA-positive myeloid cells by designing pH-selective CD28xVISTA bsAbs that act as CD28 agonists specifically within the acidic TME. These bsAbs are designed for selective “*cis*-activation” or tripartite “*trans*-activation” of CD28 in the TME, aiming for enhanced T-cell-mediated cancer cell killing while minimizing systemic T-cell activation and cytokine release syndrome (CRS) risk. *Cis*-activation relies on engagement of VISTA in the synapse formed between VISTA^+^ tumor cells or APCs and a T-cell. The *trans*-activation mechanism relies on “extra-synaptic” engagement of VISTA on myeloid cells promoting CD28 clustering on T-cells outside of the immune synapse where the initial TCR/pMHC signaling is occurring.

We report the generation and evaluation of a tumor-targeted CD28 agonist which offers co-stimulation exclusively in the presence of Signal 1 (i.e. no superagonistic properties), and potentiates the effects of a CD3-based T-cell engager (TCE; providing “artificial” Signal 1) *in vitro*. Tumor selectivity was assured by generation of a bispecific antibody monovalent for CD28 and with pH selective binding to VISTA^+^ myeloid cells in the low pH of the TME. As the Fc region of the CD28xVISTA bsAbs contains mutations abrogating cross-linking by Fcγ receptors (FcγRs), CD28xVISTA bsAbs can only induce clustering of CD28 on the surface of T-cells and hence CD28 activation when cross-linked via adjacent VISTA^+^ cells. By employing a pH-selective anti-VISTA fragment antigen–binding (Fab) domain, VISTA interaction and hence CD28 agonism is limited to the low pH of the TME with minimal risk of systemic CRS.

We show induction of CD28 signaling by CD28xVISTA bsAbs in both *cis* and *trans* in reporter cell assays *in vitro*, co-stimulation of primary human T-cell by a CD28xVISTA bsAb potentiating LNCaP prostate cancer cell killing by a CD3xPSMA T-cell engager *in vitro*, and potency in a huCD28 KI mouse model of *cis* activation. Finally, a CD28xVISTA bsAb displays good safety profile with no fortuitous induction of cytokine release or T-cell activation in physiological relevant assays.

Our approach bypasses the requirement for TAAs of a conventional CD28xTAA co-stimulatory bsAb, potentially minimizing on-target/off-tumor toxicity. Additionally, it has the capability to improve the potency of TCE’s while being cleanly combinable with any CD3xTAA and improving their safety profile/therapeutic window by orthogonally targeting of TAA and VISTA-positive cells in tumors.

## RESULTS

### Engagement of recombinant and native protein targets by CD28xVISTA bsAbs

Various formats of CD28xVISTA bsAbs were generated (Figure 1A). These were all in a human IgG1 backbone containing mutations silencing FcγR interactions (10) to avoid clustering by interaction with Fc receptors on various cell surfaces and hence minimize VISTA-independent agonism and associated CRS risks. CD28 agonism is thus completely dependent on the VISTA Fab arms engaging with their target on VISTA^+^ cells. pH-dependent agonism was achieved by incorporating VISTA binding Fab domains from clone 67375 with highly pH-selective target binding: K_D_= 0.8 nM at pH 6.0, and >400-fold lower binding at pH 7.4 (K_D_= 353 nM; (9)). Metabolism, gene and protein expression and ultimately survival of primary human T-cells is highly affected by even short term exposure to low pH *in vitro*. Therefore, a parallel set of “surrogate” bispecific antibodies were generated by incorporating VISTA Fab arms from parental antibody clone 55873 (to ensure binding to the same epitope) which bind VISTA at pH 7.4 and pH 6.0 with similar binding affinities (K_D, pH7.4_= 1 nM vs. K_D, pH6.0_= 0.6 nM, respectively; data not shown). These surrogate bsAbs allowed us to carry out cell-based *in vitro* assays at neutral pH while all *in vivo* experiments and CRS assessment studies were conducted using the pH-selective lead candidate as indicated by the notation “VISTA^pH-sens^” throughout the manuscript.

**Figure 1.**
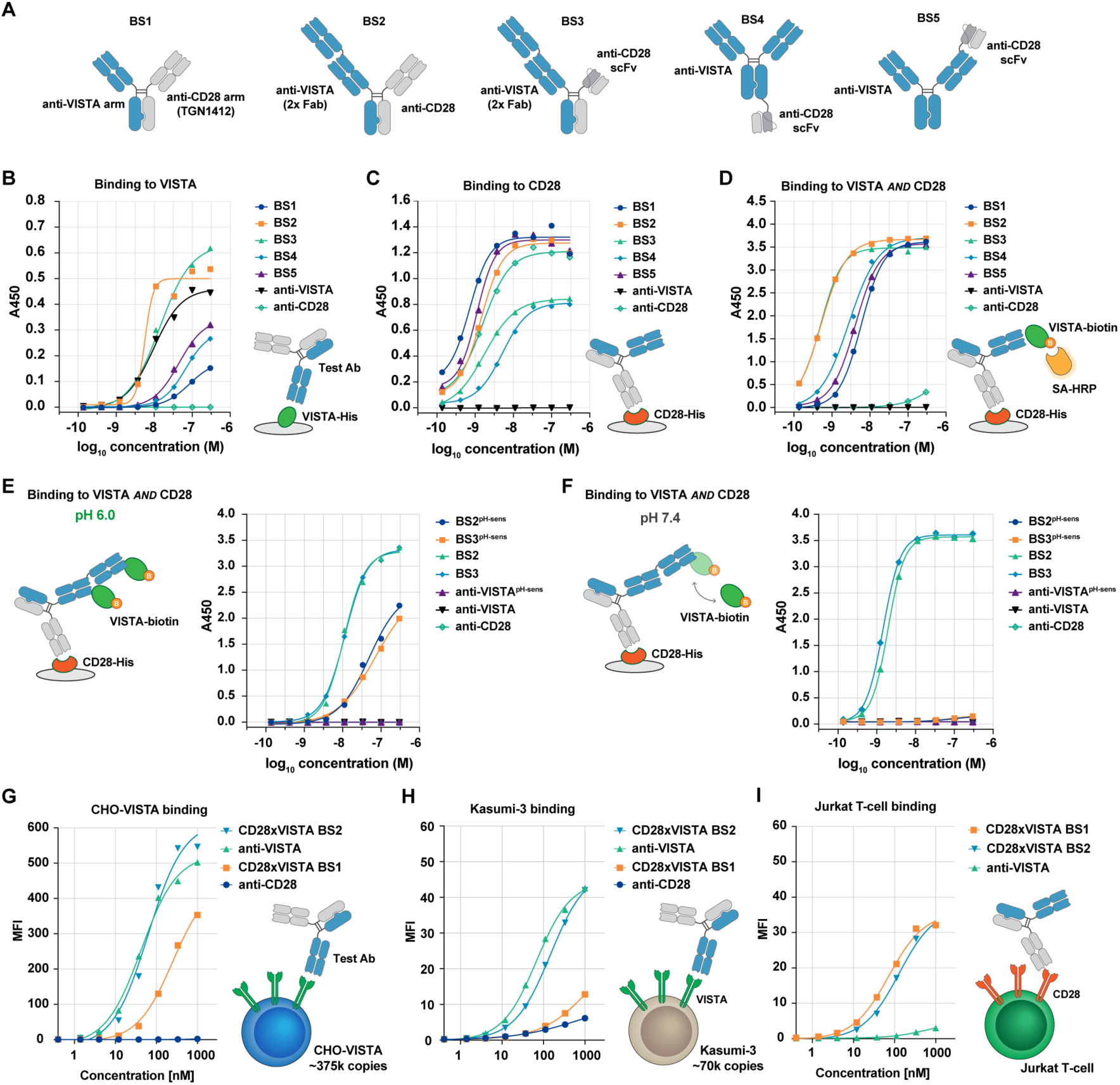
CD28xVISTA geometries BS2 & BS3 display efficient target binding. **A,** Schematic depiction of various CD28xVISTA bsAb geometries tested, with CD28-binding Fab arms or scFv domains in gray, VISTA binding Fabs in blue. VISTA binding was tested in both monovalent and bivalent formats. Both HC-LC and LC-HC orientations for the CD28-binding scFv were tested at different positions within the molecule; data for the most potent scFv configuration for each geometry is shown. **B,** Binding to recombinant VISTA protein or **C,** CD28 protein coated in ELISA wells. Parental monospecific mAbs are named anti-VISTA or anti-CD28. **D,** Simultaneous binding to VISTA and CD28 protein. **E,** Comparison of simultaneous binding by BS2 or BS3 to both targets at pH6.0 (**E**) or pH7.4 (**F**). pH-selective VISTA binding constructs are indicated by “pH-sens” superscript. **G,** Binding to native VISTA on an overexpressing CHO cell line (375.000 copies/cell) or the Kasumi-3 myeloid cell line (70.000 copies/cell) (**H**), or binding to native CD28 on Jurkat T-cells (**I**) analyzed by flow cytometry. Mean Fluorescent Intensity (MFI) was plotted as a function of mAb concentration.

We first tested binding to recombinant protein targets in an ELISA format (Figure 1B-F). Based on initial findings with our prototype BS1 format with monovalent VISTA binding, we incorporated bivalent VISTA binding in all subsequent designs (Figure 1A), which lead to improved VISTA binding efficiency through avidity (Figure 1B). All CD28xVISTA bsAb formats show dual engagement of recombinant target proteins with BS2 and BS3 formats containing duplicate VISTA binding Fab domains in a stacked arrangement connected by a flexible linker showing the most efficient simultaneous target binding in ELISA assays (Figure 1D). The mono-specific bivalent parental mAbs (from which the Fab or single-chain variable fragment (scFv) domains were derived) used as controls displayed minimal simultaneous binding in this assay (anti-VISTA and anti-CD28, Figure 1D). BS2 and BS3 formats with the pH-selective VISTA binding arm showed pH-dependent simultaneous binding to both targets, with no detectable interaction at pH 7.4 (Figure 1E & F). In parallel, the control constructs with non-selective 55873 Fab arms displayed efficient binding at both pH 6.0 and 7.4 (Figure 1E & F). The effect of avidity was further demonstrated by BS1 and BS2 bsAb binding to native VISTA on VISTA-positive cell lines with two different target densities (Figure 1G-1H), while binding to native CD28 was similar for the two formats (Figure 1I).

Taken together, CD28xVISTA bispecific formats BS2 and BS3 were equally potent in simultaneous target binding. While no issues were discovered during initial assessments, potential downstream developability issues (stability, aggregation and poor PK properties) often encountered with scFv-derived bsAbs led us to favor the BS2 format (11).

### Induction of CD28 signaling by CD28xVISTA bsAbs

Upregulation of IL-2 expression is one of the key signatures of CD28 co-stimulation (12). To assess CD28xVISTA bsAb-mediated CD28 signaling, we utilized a Jurkat-IL-2-luciferase reporter cell line. As expected from a CD28 superagonist, the addition of anti-CD28 (TGN1412) to this cell line resulted in dose-dependent expression of luciferase (Figure 2A, dark blue). In contrast, neither the prototype CD28xVISTA BS1 bsAb (Figure 2A, green) nor the monospecific parental anti-VISTA IgG1 control mAb 55873 (Figure 2A, orange) induced luciferase expression at even the highest 10 μg/mL concentration, indicating monovalent CD28 engagement does not enable CD28 clustering and signaling. In other words, our CD28xVISTA bsAb with monovalent CD28 binding did not display superagonism in this experimental setting.

**Figure 2.**
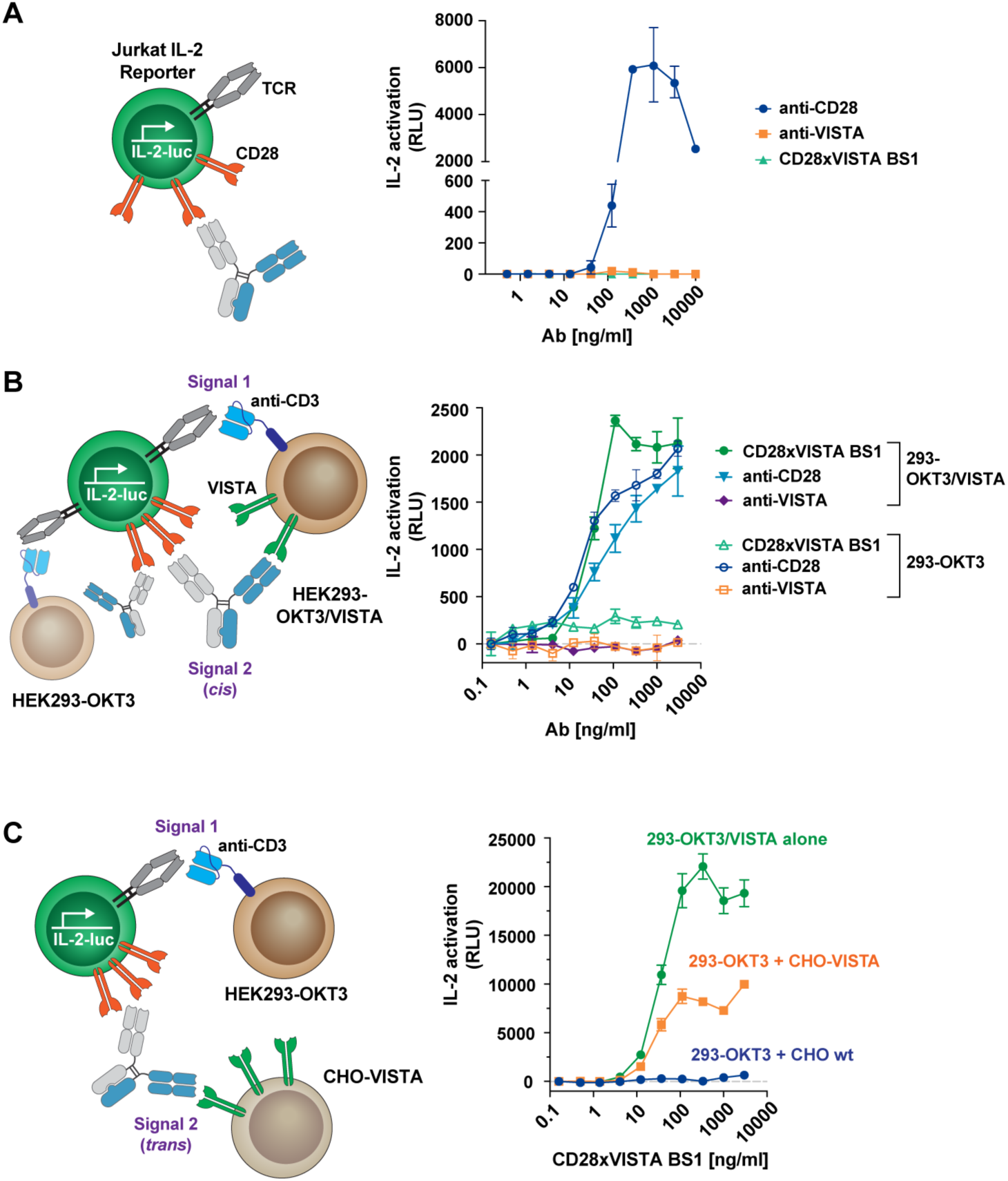
CD28xVISTA bsAb induces IL-2–luciferase reporter gene expression both in *cis* and in *trans*. **A,** Luciferase expression from a Jurkat IL-2-luciferase reporter cell line as a function of concentration of CD28xVISTA BS1 (green) or parental monospecific mAbs anti-VISTA (orange) or anti-CD28 (blue). **B,** Luciferase expression from the reporter cell line co-cultured with a HEK293 cell line expressing anti-CD3 (OKT3-scFv) alone (open symbols; left in schematic) or in combination with VISTA (filled symbols; right in schematic) enabling co-stimulation by a CD28xVISTA bsAb in *cis*. **C,** Luciferase expression from the IL-2 reporter cell line as a function of CD28xVISTA BS1 when co-cultured with the HEK293 cell line expressing anti-CD3 (OKT3-scFv) in combination with a CHO cell line overexpressing VISTA (orange) enabling co-stimulation by a CD28xVISTA bsAb in *trans*. A parallel negative control with the parental CHO cells substituting for CHO-VISTA cells (blue) or a positive control with the HEK293 cell line co-expressing anti-CD3 (OKT3-scFv) and VISTA (allowing for *cis* co-stimulation; green) are indicated.

In contrast, when the IL-2 reporter cell line was co-cultured with HEK293 cells co-expressing membrane-anchored anti-CD3 scFv (OKT3-scFv) and VISTA, a robust dose-response effect on luciferase expression was seen with BS1 in this *cis*-activation setting (Figure 2B, dark green). Luciferase expression was in fact somewhat higher than seen with the TGN1412 control (Figure 2B, light blue). A parallel control experiment with the Jurkat reporter cell line co-cultured with HEK293 cells expressing only OKT3-scFv (without VISTA) lead to only minimal luciferase expression by BS1 (Figure 2B, light green), highlighting that monovalent CD28 engagement by BS1 without simultaneous binding to VISTA does not lead to CD28 signaling. In both settings, the monospecific parental anti-VISTA IgG1 control mAb 55873 led to only background luciferase expression levels (Figure 2B, purple, orange).

To investigate whether our prototype CD28xVISTA bsAb BS1 could potentially induce *trans*-activation (i.e. Signal 2 provided by interaction between T-cell and a cell different from the Signal 1-inducing cell line), the Jurkat reporter line was co-cultured with the HEK293 cells expressing only OKT3-scFv (no VISTA) as well as CHO-K1 cells engineered to overexpress VISTA. While luciferase expression from the Jurkat cells was not induced by BS1 when co-cultured with the 293 OKT3 cell line and CHO cells not expressing VISTA (Figure 2C, blue), inclusion of CHO cells expressing VISTA enabled a robust induction of luciferase expression by addition of CD28xVISTA BS1 (Figure 2C, orange), albeit not as strong as seen when OKT3 and VISTA was co-expressed on the same HEK293 cells, i.e. *cis* activation (Figure 2C, green).

In conclusion, the prototype CD28xVISTA BS1 bsAb induced VISTA-dependent IL-2-luciferase reporter expression in *cis* as well as in *trans*.

### Human T-cell co-stimulation by CD28xVISTA BS2 potentiates LNCaP killing by a CD3xPSMA T-cell engager

To characterize the co-stimulatory activity of a CD28xVISTA bsAb in a physiologically more relevant setting, we tested the effect of CD28xVISTA BS2 on primary human T-cell-mediated killing of LNCaP prostate cancer cells *in vitro*. Co-cultures of LNCaP cells with human PBMCs and the VISTA-positive Kasumi-3 myeloid cell line in the presence of a CD3xPSMA TCE alone or in combination with CD28xVISTA BS2 were analyzed on the xCELLigence real-time cell analysis platform (Figure 3A). The use of a TCE allowed for a facile way to fine-tune TCR stimulation (“Signal 1”) to explore the co-stimulatory effect over a broad range of sub-maximal TCR stimulation levels. LNCaP cells do not express VISTA (Supplementary Figure 1), so a potential effect on cell killing would not be mediated by a *cis* co-stimulatory effect (Signal 1 and Signal 2 both occurring simultaneously in the synapse between a T-cell and LNCaP cell). Inherent to the design of the assay platform, only growth of the adherent LNCaP cells caused a change in the measured impedance signal; no effect from non-adherent PBMC’s or Kasumi-3 cells was detected.

**Figure 3.**
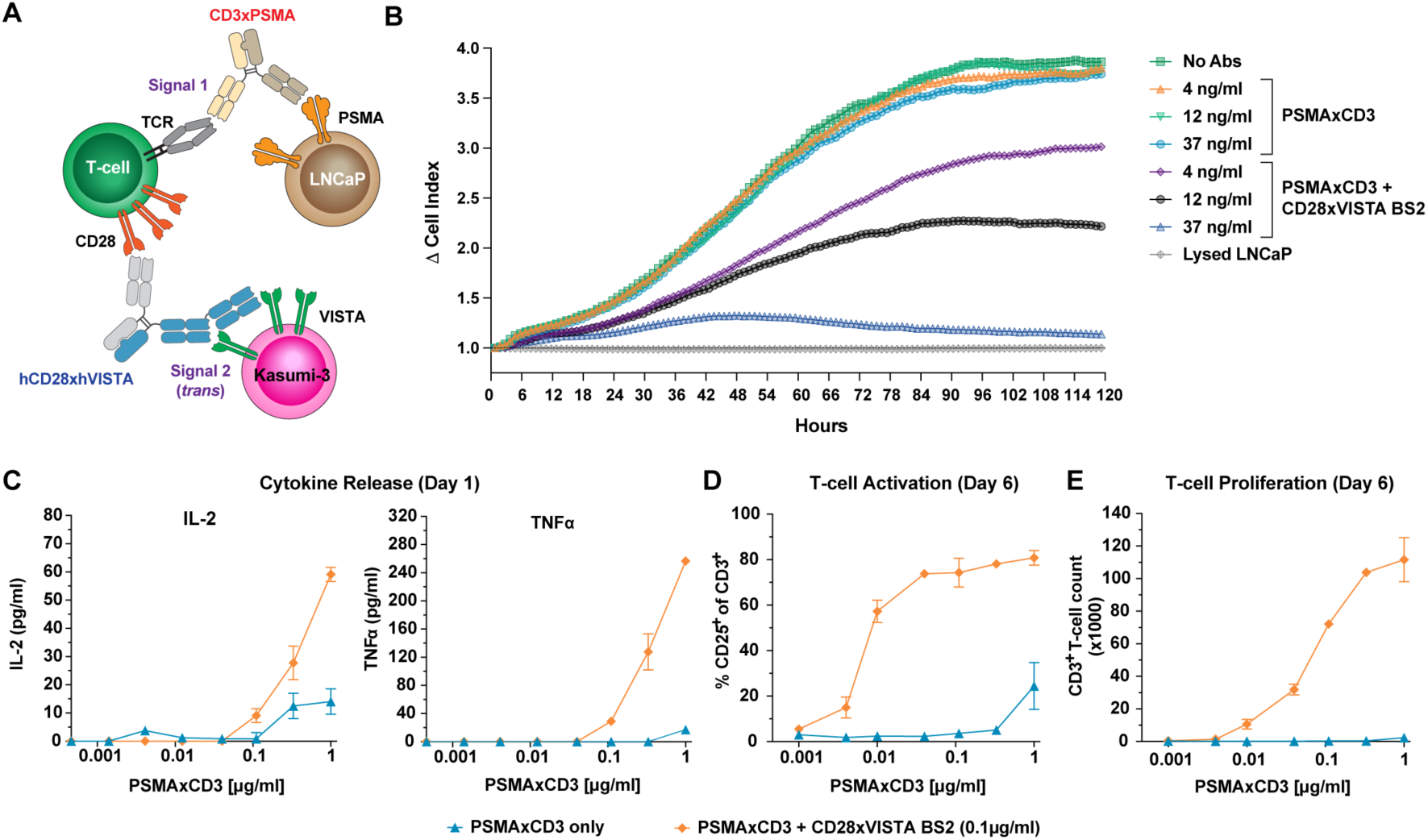
*In vitro* human T-cell co-stimulation in *trans* by CD28xVISTA BS2 (non-pH sensitive) potentiates LNCaP killing by a CD3xPSMA T-cell engager. **A,** Schematic representation of LNCaP prostate cancer cells co-cultured with Kasumi-3 and T-cells from human PBMCs. LNCaP killing by a CD3xPSMA TCE can potentially be enhanced through CD28 *trans* co-stimulation by CD28xVISTA BS2 **B,** Signal resulting from LNCaP cells in the Agilent xCELLigence assay platform in the absence of mAbs (green squares), in the presence of increasing sub-optimal concentrations of CD3xPSMA alone (4 ng/ml (orange), 12 ng/ml (light green) or 37 ng/ml (blue)); or same concentrations of CD3xPSMA in the presence of 0.1 μg/ml CD28xVISTA BS2 (purple, black, and blue triangles, respectively). A control with lysis buffer added to co-culture wells is indicated by gray. Only growth of the adherent LNCaP cells caused a change in the impedance signal measured by the xCELLigence platform. **C,** IL-2 or TNF-α cytokine release measured Day 1, (**D**) T-cell activation measured as %CD25^+^ of total T-cells on Day 6 and (**E**) T-cell proliferation enumerated as T-cell count per volume on Day 6 in the absence (blue) or presence of 0.1 μg/ml CD28xVISTA BS2 (orange) (mean values +/-SD; n=3 technical replicates).

LNCaPs cultured with Kasumi-3 and T-cells with no added test antibodies reach confluence in approximately 5 days (120 hr; Figure 3B, dark green). Inclusion of CD3xPSMA at ∼4, 10 and 40 ng/mL did not substantially affect growth of the adherent LNCaP cells compared to co-cultures in the absence of the TCE (Figure 3B). However, those same concentrations lead to substantial dose-dependent decrease of LNCaP viability in the presence of 100 ng/mL CD28xVISTA BS2, with the highest 40 ng/mL TCE concentration leading to complete killing of LNCaP cells as indicated by a signal at day 5 comparable to wells with addition of lysis buffer (Figure 3B). The potentiation of LNCaP killing by the CD3xPSMA TCE was consistent across 3 different donors (data not shown).

Culture supernatant samples retrieved from the plate after 1 day of bsAb treatment were analyzed for cytokine induction, and T-cell activation (upregulation of CD25 expression on T-cells, a commonly used activation marker responsive to CD28 co-stimulation) and proliferation (CD3^+^ T-cell count) were measured in samples retrieved after 6 days using flow cytometry. Enhanced killing in the presence of the CD28xVISTA bispecific antibody was accompanied by increased IL-2 and TNF-α cytokine release (Figure 3C), T-cell activation (Figure 3D) and proliferation (Figure 3E). No effect of the presence of CD28xVISTA BS2 on any of these readouts was observed in the absence of CD3xPSMA, demonstrating that a TCR signal is required to enable the CD28 co-stimulatory effect. No superagonistic properties of CD28xVISTA BS2 were seen at the 0.1 µg/mL concentration used in this experiment.

### CD28xVISTA BS2 inhibits MC38-hVISTA tumor growth in hCD28 KI mice in combination with anti-PD-1

Tumor growth inhibition (TGI) by CD28xVISTA^pH-sens^ BS2 (with pH-selective hVISTA engagement) of MC38 cells overexpressing human VISTA was tested in a humanized CD28 syngeneic mouse model alone or in combination with anti-murine PD-1 (anti-mPD-1). In this model, tumor-specific T-cells were activated by their TCRs engaging peptide/MHC complexes on the tumor cells, i.e. a natural “Signal 1” which could potentially be enhanced by CD28 co-stimulation in *cis* between T-cells and hVISTA^+^ tumor cells (Figure 4A).

**Figure 4.**
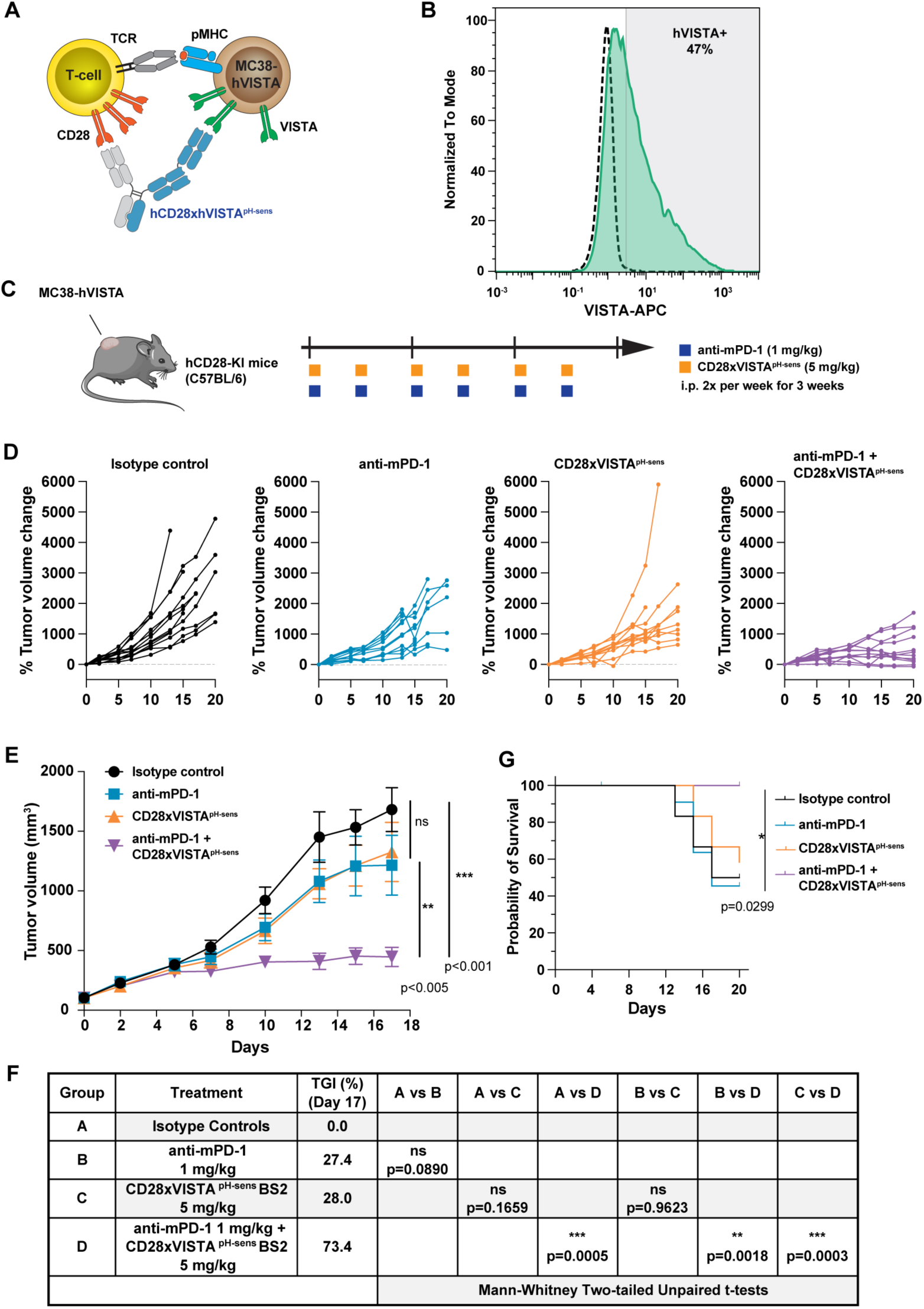
pH selective CD28xVISTA^pH-sens^ BS2 bsAb inhibits MC38-hVISTA tumor growth in hCD28 KI mice in combination with anti-PD-1. **A,** Schematic of T-cells from the hCD28-KI mice expressing human CD28 receiving CD28 co-stimulation in *cis* through CD28xVISTA binding to hVISTA^+^ MC38 cells. **B,** Distribution of hVISTA^+^ MC38 cells prior to implantation as measured by flow cytometry with a fluorescently labelled anti-hVISTA mAb. Isotype control staining shown by black dotted line. **C,** Schematic of dosing schedule following subcutaneous injection of a population of 1 × 10^6^ cells according to (**B**) into hCD28 KI mice and randomization into 4 groups (n=12/group) dosed ip with the indicated doses of either isotype controls, anti-mPD-1 (blue), CD28xVISTA^pH-sens^ BS2 (orange), or anti-PD-1 + CD28xVISTA^pH-sens^ BS2. **D,** Spider plots of tumor volume measurements for individual mice in each treatment group: isotype controls (black), anti-mPD-1 (blue), CD28xVISTA^pH-sens^ BS2 (orange), or anti-PD-1 + CD28xVISTA^pH-sens^ BS2 (purple). **E,** Tumor volume per group represented as mean values +/- SEM; color coding as in (**D**). **F,** Tumor growth inhibition (TGI) and statistical significance of endpoint MC38 tumor volumes evaluated using the Mann-Whitney two-sided unpaired t test with exact *P* values (*0.01 < *P* < 0.05; ** 0.001 < *P* < 0.01; *** 0.0001 < *P* < 0.001). **G,** Survival curves, with significance determined using the Mantel-Cox log-rank test (\**P* = 0.0299). Color coding as in (**D**). Tumor growth inhibition calculations and significance criteria are as described (13).

MC38 cells overexpressing human VISTA were used for implantation into female human CD28 knock-in (KI) mice. Expression of hVISTA on the preconditioned cells prior to implantation was analyzed by flow cytometry using a fluorescently labelled anti-human VISTA mAb. As indicated by the overlap of staining with an isotype control mAb, a substantial population of MC38 cells were not expressing hVISTA (Figure 4B). Once the tumor volumes reached ∼80-100 mm^3^ (day 7 after implantation) mice were randomized into 4 groups of 12 animals per group. Animals were administered intraperitoneally twice/week with isotype controls, a suboptimal dose of 1 mg/kg anti-mPD-1, CD28xVISTA^pH-sens^ BS2 at 5 mg/kg, or anti-mPD-1 (1 mg/kg) + CD28xVISTA^pH-sens^ BS2 (5 mg/kg) combined (Figure 4C). A modest, non-significant effect on tumor growth of anti-mPD-1 or CD28xVISTA^pH-sens^ BS2 monotherapy of 27.4%, and 28.0%, respectively, was seen compared to the isotype control arm (Figure 4E & F). However, combining CD28xVISTA^pH-sens^ BS2 with the low dose anti-PD-1 resulted in a tumor growth inhibition of 73.4% on day 17, a significant reduction both compared to the isotype control group (*P*= 0.0005) and either of the monotherapy arms (*P*= 0.0018; Figure 4E & F). This effect translated to a significant increase in probability of survival between combination arm and control group (*P*= 0.0299; Figure 4G). Interestingly, the substantial tumor growth inhibition and enhanced survival was observed despite the highly heterogeneous tumor cell population (of the MC38 cells inoculated only 47% were hVISTA^+^), suggesting efficient tumor growth control of even MC38 cells not expressing hVISTA. Importantly, as the anti-VISTA Fab arms of CD28xVISTA^pH-sens^ BS2 are not cross-reactive to murine VISTA, the effect on tumor growth can be uniquely ascribed to the CD28 agonistic effect, while potential VISTA checkpoint inhibition does not play a role.

In conclusion, CD28xVISTA^pH-sens^ BS2 enhanced natural “Signal 1” by CD28 co-stimulation in *cis* resulting in significant TGI in combination with anti-mPD1.

### Pharmacokinetic profile of CD28xVISTA BS2 bsAb in huCD28 KI mice is not impacted by high affinity CD28 binding

CD28 expressed on T-cells in blood and lymph nodes could provide a significant antigen sink. We wanted to explore a potential effect of CD28 binding affinity on the PK properties of a CD28-targeting bsAb. We compared the pharmacokinetics of 3 highly similar bsAbs in huCD28 KI mice, each having a different CD28 binding domain with binding affinities spanning two orders of magnitude (K_D_ values of 1-2 nM for CD28xVISTA^pH-sens^ BS2 (SPR (14,15) to 272 nM for R-5678 (SPR; 37°C (16)). CD28^ADI^xVISTA^pH-sens^ BS2 has an identical architecture to CD28xVISTA^pH-sens^ BS2 except for a different anti-CD28 Fab arm with approx. 60-fold lower affinity compared to the CD28 binding arm of CD28xVISTA^pH-sens^ BS2 (data not shown). R-5678 is a CD28xPSMA bsAb using the same Fc region as CD28xVISTA^pH-sens^ BS2. The PSMA arm of R-5678 is not cross-reactive to murine PSMA.

When dosed IV at 5 mg/kg in human CD28 knock-in mice, which systemically expresses human CD28, all 3 antibodies showed relatively slow clearances with differences that did not correlate with CD28 binding affinity (Figure 5), indicating that other intrinsic factors besides CD28 affinity are dominant in terms of resulting PK. We have no explanation for the low C_max_ translating into lower AUC/exposure values for R-5678. The calculated clearance rate of CD28xVISTA^pH-sens^ BS2 falls in the middle of the range found for 16 human non-mouse cross-reactive IgG1 mAbs (17), and is similar to the parental monospecific, bivalent pH-selective anti-VISTA IgG1 mAb with the same anti-VISTA Fab arms dosed at the same level and route in a wt C57BL6/J mouse (9). The pH-selective VISTA Fab arms employed in CD28xVISTA^pH-sens^ BS2 in the context of a regular bivalent IgG1 mAb format showed favorable PK and did not provide TMDD in hVISTA KI mice, cynomolgus monkeys or human trial subjects (SNS-101(9,18)). However, as VISTA binding is not mouse cross-reactive, any negative effect on PK in the bispecific context would not be revealed in this hCD28 KI mouse model.

**Figure 5.**
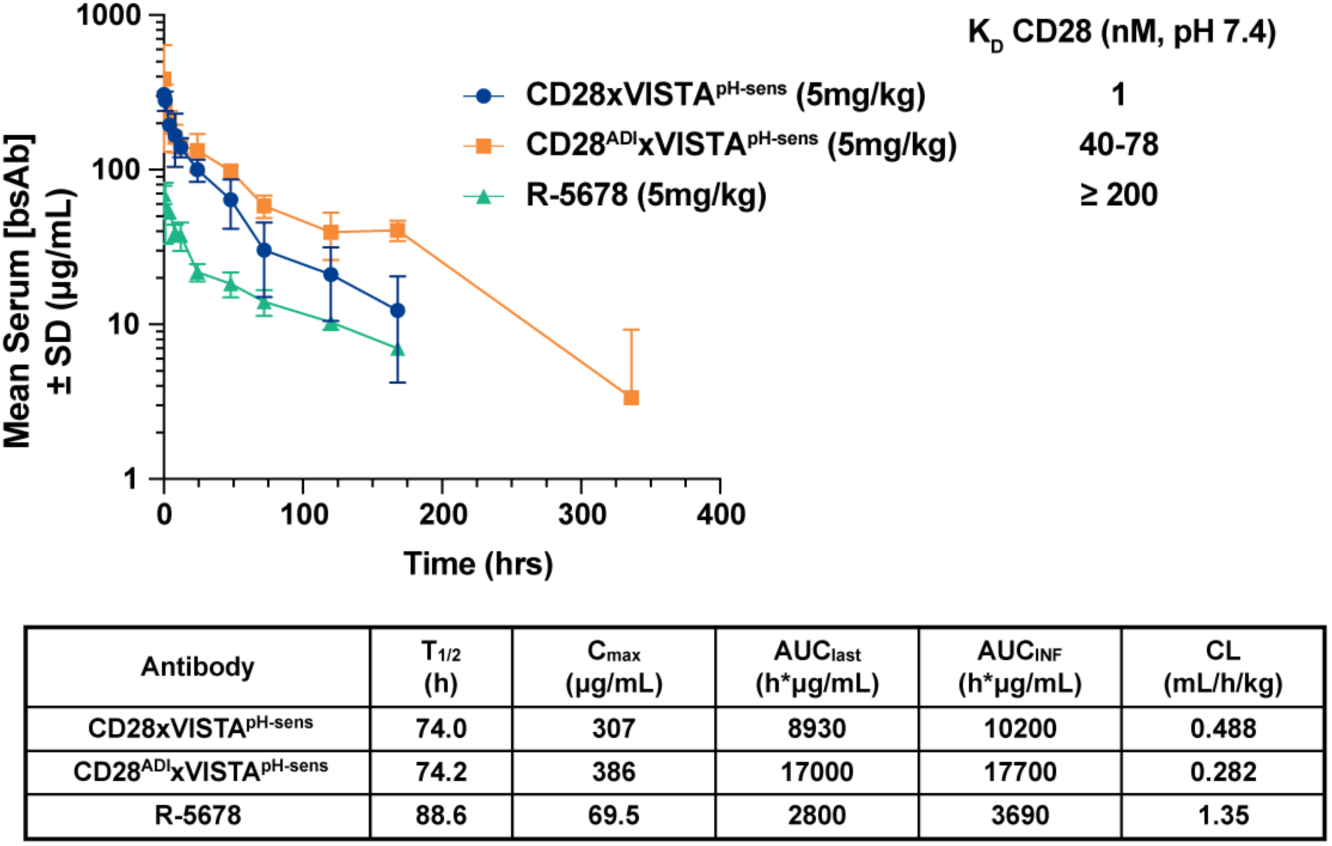
PK profile of CD28xVISTA^pH-sens^ BS2 in huCD28 KI mice. Human CD28 KI mice (n=4 per group) were given a single intravenous injection of 5 mg/kg CD28xVISTA^pH-sens^ BS2 (blue) or comparators CD28^ADI^xVISTA^pH-sens^ BS2 (orange) or R-5678 (green). Serum samples were collected over a 28-day observation period and levels of bsAb were determined by ELISA. Data are presented as mean values +/- SD. Calculated PK parameters are shown in the inset table. K_D_ values collected by Octet BLI (BS2) or SPR (R-5678).

Our data suggests a favorable PK profile of CD28xVISTA^pH-sens^ BS2 in huCD28 KI mice, and that the low K_D_ of the CD28 binding domain in a monovalent format did not result in unacceptable rapid clearance.

### CD28xVISTA displays good safety profile with no induction of cytokine release in physiological relevant assays

We conducted cytokine release assays developed to predict the CD28 superagonistic activities of therapeutics in pre-clinical development. We first assessed the CRS potential of CD28xVISTA^pH-sens^ BS2 in co-cultures of human umbilical vein endothelial cells (HUVECs) and human PBMCs (19). In this assay, CD28xVISTA^pH-sens^ BS2 was compared to the previously described bsAbs CD28^ADI^xVISTA^pH-sens^ BS2, R-5678, as well as CD28 superagonist TGN1412 as a positive control. Antibody concentrations of 1, 10, 30 and 100 µg/ml were incubated in the co-cultures for 48h prior to cytokine analysis. TGN1412 resulted in the highest level of induction of all 8 cytokines tested, while CD28xVISTA^pH-sens^ BS2 showed only minimal propensity for cytokine release (Figure 6A). CD28^ADI^xVISTA^pH-sens^ BS2 induced a slightly increased response compared to CD28xVISTA^pH-sens^ BS2, particularly for IFN-γ, IL-2 and IL-6. Interestingly, the comparator R-5678 with Fab arms from Regeneron’ s clinical CD28xPSMA IgG4 candidate Nezastomig placed in the IgG1 Fcγ null backbone used in CD28xVISTA^pH-sens^ BS2 did display cytokine release in multiple donors at higher concentrations, particularly of IFN-γ, IL-4, IL-6 and IL-8. This could be a result of low level PSMA expression on HUVECs.

**Figure 6.**
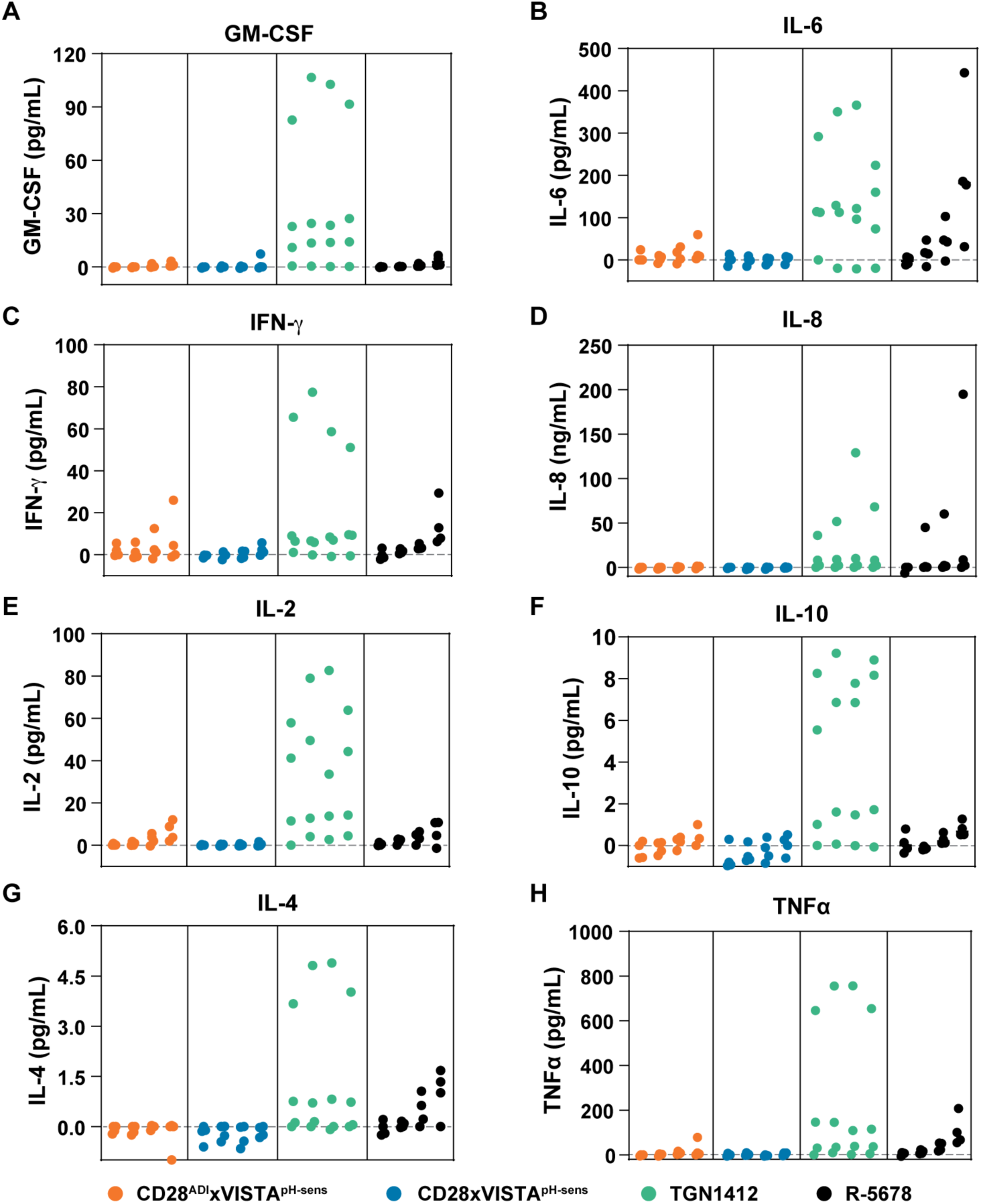
*In vitro* CRS assessment in HUVEC:PBMC co-culture assays. Levels of cytokines GM-CSF, IL-6, IFN-γ, IL-8, IL-2, IL-10, IL-4, and TNFα (**A, B, C, D, E, F, G,** and **H,** respectively) were measured 2 days after incubation with CD28^ADI^xVISTA^pH-sens^ BS2 (orange), CD28xVISTA^pH-sens^ BS2 (blue), TGN1412 (green) or R-5678 (black) at concentrations of 1, 10, 30, and 100 μg/ml. Quantification was done with 6 replicates for each concentration of each mAb. Each dot represents the results from one human donor (4 donors tested).

In a second assay, a cytokine release profile of CD28xVISTA^pH-sens^ BS2 was generated in an *ex vivo* platform that mimics human blood circulation for a more accurate CRS prediction (Figure 7A; (20)). *Ex vivo* cytokine release in fresh human whole blood from 6 healthy donors by CD28xVISTA^pH-sens^ BS2 was compared to CD28^ADI^xVISTA^pH-sens^ BS2 and R-5678 at concentrations of 1, 10 and 100 μg/ml (mirroring realistic clinical plasma levels) and the controls anti-CD28 (ANC.28.1; 1 μg/ml), Alemtuzumab (3 μg/ml) or Cetuximab (250 μg/ml). These controls exhibited the expected effects on cytokine release (Figure 7B-F) while none of the tested bispecific Abs induced cytokine release significantly different from PBS or formulation buffer up to and including the highest tested concentration of 100 μg/mL, an estimated peak serum concentration reached by a 5 mg/kg IV dose of the pH-selective anti-VISTA IgG1 mAb SNS-101 in humans (18).

**Figure 7.**
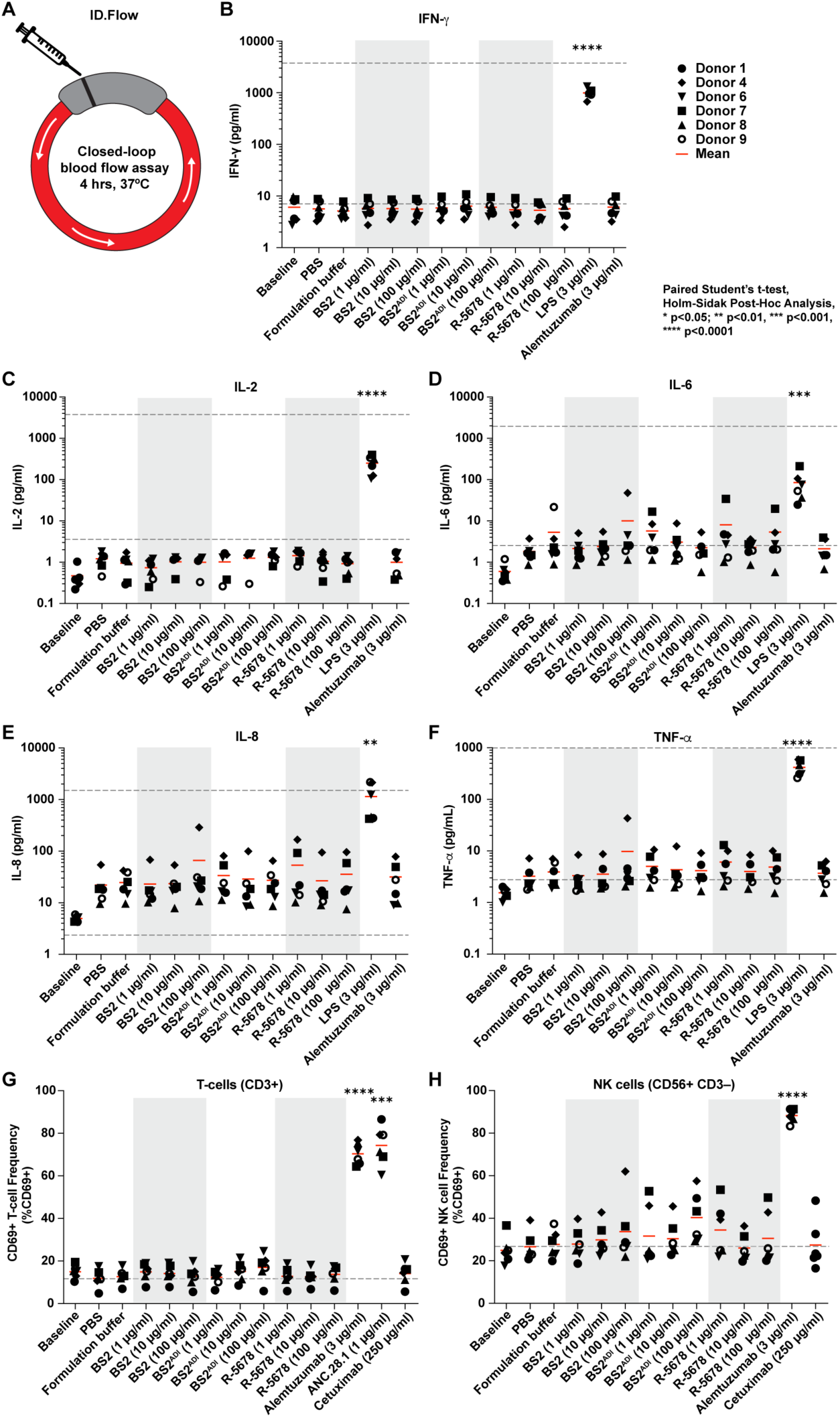
pH selective CD28xVISTA^pH-sens^ BS2 does not induce cytokine responses, T-cell or NK-cell activation in sensitive *ex vivo* human whole blood ID.Flow assay. **A,** Schematic representation of the closed-loop ID.Flow assay platform. **B,** through **F,** measured levels of IFN-γ, IL-2, IL-8, IL-6, and TNFα, respectively, after incubation for 4h with the antibodies and concentrations indicated in freshly collected, circulating blood from 6 healthy human volunteers. **G,** Activated T-cells and (**H**) NK cells gated as CD69^+^ cells and presented as the percentage of total cells of the respective cell type as measured by flow cytometry. Appropriate positive controls with known effects on the test parameters were included, i.e., alemtuzumab (anti-CD52), and anti-CD28 (ANC28.1). Phosphate buffered saline (PBS), cetuximab (anti-epidermal growth factor receptor [EGFR] antibody) and formulation buffer for the 3 tested bsAbs were used as negative controls. BS2: CD28xVISTA^pH-sens^; BS2^ADI^: CD28^ADI^xVISTA^pH-sens^; R-5678: CD28xPSMA. Lower level of quantification indicated by dotted lines; mean values in red line. A single n= 1 experiment was performed with n= 6 independent donor samples. *p<0.05; **p<0.01; ***p<0.001; ****p<0.0001 comparison to formulation buffer (**B** through **H**) by Paired Student’s t-test with Holm-Sidak correction.

In addition, no significant effect on T-cell (**G**) or NK cell activation (**H**) was observed. No effect on blood cell viability including platelet (PLT), white blood cell (WBC), and red blood cell (RBC) was found (Supplementary Fig. 2), and in agreement with the lack of cytokine release, granulocyte and monocyte counts and activation, as well as T and NK cell counts were not significantly affected by any of the test items at all three concentrations (Supplementary Fig. 2).

In conclusion, a favorable safety profile of CD28xVISTA^pH-sens^ BS2 in two different *in vitro* assays for CRS risk assessment was observed. We consider these assays more predictive of potential CRS issues in humans than NHP toxicity studies given the past failure of NHPs to predict the negative effects of the CD28 superagonist TGN1412 (21).

## DISCUSSION

Agonism of positive T-cell co-stimulatory signals in the context of existing checkpoint inhibitors, T-cell engagers, or CAR T-cells is a relatively new therapeutic avenue to boost antitumoral efficacy. Specifically, interest in tumor-targeted CD28 bsAbs has been growing in recent years after safety concerns had halted development of CD28 agonists for over a decade after the failed TGN1412 clinical trial (5). Recent evidence suggest that many “warm” and “hot” tumors have resident anti-tumor T-cells with the capacity to kill malignant cells (22–26). However, these T-cells are often functionally “exhausted” but may be reawakened by co-stimulation through CD28 activation.

In the potency study in hCD28 KI mice described here, CD28xVISTA^pH-sens^ BS2 significantly inhibits MC38-hVISTA tumor growth inhibition in combination with anti-PD-1. The immunity of the MC38 tumor cells afforded a natural Signal 1 and CD28 co-stimulation in *cis* provided enhanced tumor control. Reports suggest PD-1 suppresses T-cell function primarily by inactivating CD28 signaling (CD28 is the main target for dephosphorylation by PD-1-recruited Shp2 phosphatase in response to PD-1 activation by PD-L1 (27)), so a combinatorial effect of anti-PD1 and CD28 co-stimulatory bsAbs is expected. This positive effect has been observed in other preclinical studies (e.g. (3)). Furthermore, CD28 signaling was necessary for restoring T-cell responses during mAb PD-1 blockade in tumor-bearing mice (28).

A lack of cross-reactivity to mouse VISTA means *cis* co-stimulation is the only way to explain the potency of CD28xVISTA^pH-sens^ BS2 in the hCD28-KI study as VISTA blockade itself cannot occur and contribute to the observed efficacy. To make the experiment more physiologically relevant, we utilized a population of cells with highly variable hVISTA expression (with about half of MC38 cells showing a hVISTA signal similar to isotype control staining by flow cytometry) instead of implanting a homogeneous clone highly overexpressing hVISTA. Half of hVISTA^+^ MC38 cells seem to provide enough co-stimulation to generate a broader effect on TGI. We envision that bystander killing of MC38 cells not expressing hVISTA could be explained by a general expansion in tumor-reactive T-cells. This result could bode well for addressing the issue of “real-world” patient tumor heterogeneity.

Analysis of the TCGA Pan-Cancer Atlas RNA-seq dataset controlling for leukocyte infiltration in samples identified a subset of cancers with VISTA expression directly on tumor cells (29). In addition, immunohistochemistry experiments have suggested VISTA expression on human pancreatic ductal adenocarcinoma, ovarian and endometrial cancer cells (30,31). However, VISTA expression in malignant cells is generally very low, particularly relative to myeloid cell types like macrophages and dendritic cells, as evidenced by pan-cancer scRNA-seq atlases (32). Cancers like these with appreciable VISTA expression on tumor cells may define a group where *cis*-activation by a CD28xVISTA bsAb is sufficient for inhibiting tumor growth, and could be explored further with respect to patient stratification. Importantly, T-cell co-stimulation by a CD28xVISTA bsAb appears to also work independently of VISTA expression directly on tumor cells as described below.

As previously mentioned, VISTA is highly expressed on human hematopoietic cells and at the highest densities in the myeloid lineage (monocytes, macrophages, dendritic cells (DCs)) based on flow cytometry. Often considered the most potent antigen presenting cells (APCs), DCs are crucial for initiating and regulating immune responses. A two-step model of CD8^+^ T-cell activation was recently proposed: Initial activation and proliferation of tumor-specific CD8^+^ T-cells in tumor-draining lymph nodes maintaining a stem-cell-like phenotype, and subsequent effector program acquisition within the tumor requiring CD28 co-stimulation provided by CD80- and CD86- expressing mature DCs (33). This model highlights CD28 co-stimulation as a critical signal within the TME that regulates anti-tumor CD8^+^ T-cell differentiation and acquisition of an effector phenotype. We postulate that the interaction of CD80 and CD86 on mature DCs with CD28 on T-cells ensuring their full activation could be replaced or augmented with a CD28xVISTA bsAb bridging the mature VISTA^+^ DC and T-cell, providing CD28 co-stimulation in *cis*. This setting is mimicked in the IL2-reporter assay in Figure 2B where the 293 cell line co-expressing membrane anchored anti-CD3 scFv as well as VISTA can be considered an “artificial” DC. There is further evidence for “immune triads” between APCs, antigen-specific CD4^+^ and CD8^+^ T-cells that are potent sites of instruction for anti-tumor CD8^+^ T-cells (34,35), and we speculate that an APC-engaging CD28xVISTA bispecific could potentially enhance the stability and potency of such immune triads.

In sum, CD28-mediated co-stimulation of T-cells in *cis* by a CD28xVISTA bsAb could occur both through VISTA^+^ tumor cells in the TME as well as through VISTA^+^ antigen-presenting mature DCs in tumors or lymph nodes; both environments characterized by low pH.

Our Jurkat IL-2-reporter cell assays indicate a CD28xVISTA bsAb can provide CD28 activation not only in *cis* (between T-cell and cell displaying p/MHC complex) but also in *trans* (with stimulation occurring through clustering mediated by a third VISTA^+^ cell outside of the synapse). We also show that CD28xVISTA BS2 potentiates LNCaP cancer cell killing by a CD3xPSMA TCE *in vitro*. This potentiation could only happen through *trans*-activation as LNCap’s do not express VISTA (Supplementary Figure 1). Likely, CD28xVISTA BS2 combined with the CD3xPSMA TCE enhance target cell killing through IL-2-dependent expansion of CD25^+^ T-cells. As proposed for a TAA-targeted CD28 bsAb, induction of IL-2 secretion along with upregulation of the IL-2 receptor (CD25) could potentially create an autocrine positive feedback loop for tumor-reactive T-cells (36). A shift towards a more favorable Effector:Target cell ratio by CD28 co-stimulation-mediated proliferation would likewise be highly beneficial in patients, turning “lukewarm” tumors “hot”. We relied on a TCE as a tool to provide artificial Signal 1 in an easily tunable fashion, however we expect the *trans* co-stimulation effect to apply to not only combination with a TCE but also to settings with natural Signal 1 as replicated in our *in vivo* efficacy study in hCD28 KI mice.

While traditionally *cis* co-stimulation was reported to be much more potent than *trans* for both CD28 (37) as well as 4-1BB (38), several reports of CD28 *trans* co-stimulation have been published (39–42). Additionally, a *trans*-activation concept has been utilized for a 4-1BB agonist in the form of tumor stroma antigen FAPx4-1BB bispecific constructs (4). Such a bsAb has progressed to clinical trials (43). Several difficult to treat tumor types, including hepatocellular carcinoma, pancreatic adenocarcinoma, and glioblastoma, have extremely high myeloid content, to the extent that the myeloid cells make up the bulk of the tumor mass (44–46). A high proportion of VISTA^+^ myeloid cells and frequent myeloid:T-cell interactions would provide ample opportunities for a bispecific myeloid-T-cell targeting bsAb to act. We are currently developing a hCD28xhVISTA double knock-in mouse model that will be a prerequisite for *in vivo* testing of the myeloid cell engagement setting in *cis* or *trans*.

Minimizing systemic activation is crucial for the generation of a safe CD28 agonist. All other disclosed bsAb programs targeting CD28 rely on engagement of a TAA (5). Highly specific antigens for solid tumors are extremely rare, only antigens with differential expression on tumor vs. normal cells exist. This leads to on-target, off-tumor toxicity and narrow therapeutic windows. Conditionally active mAb technologies have previously been applied to T-cell co-stimulatory antibodies to restrict activity to the tumor. ATP-dependent binding moieties (47) as well as conditional steric blockade approaches (48) have been employed in 4-1BB agonists. Our novel pH-selective targeting approach generates a tumor-specific bispecific CD28 agonist while avoiding engagement of TAAs of ambiguous selectivity.

VISTA^+^ myeloid cells are abundant not only in the TME but also outside the tumor. We previously described how pH-selective engagement of VISTA by a bivalent IgG1 format abolished binding to VISTA^+^ cells at neutral pH, avoiding TMDD and CRS issues associated with non-pH selective anti-VISTA mAbs (9). While high affinity CD28 binding could potentially affect PK of a CD28xVISTA bsAb, our PK study in hCD28 KI mice did not reveal a correlation between CD28 binding affinities and PK properties in a monovalent binding format. The lack of systemic binding to VISTA^+^ cells by our CD28xVISTA^pH-sens^ BS2 bsAb at neutral pH was reflected in our safety assessment (HUVEC:PBMC co-culture assays as well as *ex vivo* human blood circulation assay) suggesting a very low propensity for systemic cytokine induction and T-cell activation. Incorporating FcR null mutations into TGN1412 itself abolishes its superagonistic activity (15,49), and the monovalent CD28 engagement in addition to the FcR null mutations in our CD28xVISTA bsAb designs further reduces the risk of fortuitous CD28 clustering and activation in the absence of VISTA engagement.

Extra care must be taken when developing CD28 agonists, as superagonism must not only be avoided as illustrated by the TGN1412 clinical trial (7), but additional attributes which cannot easily be assessed in *ex vivo* assays or in animal models need to be carefully considered. In clinical trials in cancer patients with the CD28xPSMA bispecific co-stimulatory antibody REGN5678 (Nezastomig) as well as an Fc-fusion of an engineered CD80 variant with increased affinity for PD-L1 and CD28, ALPN-202 (Davoceticept) (50), combinations with PD-1 blockade showed meaningful clinical benefits but also revealed a significant safety risk (51,52). The REGN5678 and cemiplimab combination was associated with two immune-related deaths in a trial of advanced prostate cancer that resulted in a study halt and subsequently refocused efforts to testing REGN5678 either as monotherapy, in combination with decreased doses of cemiplimab, or alternatively a CD3xPSMA T-cell engager(53). Importantly, another TAA targeted by REGN7075 (CD28xEGFR) did not show similar problems in combination treatment with cemiplimab (53), indicating the toxicity profile in REGN5678 could be due to on-target/off-tumor effects as PSMA is known to be expressed on normal tissue (54), and not necessarily a CD28 agonist drug class effect. Our candidate showed a benign safety profile in cytokine release and T-cell activation assays with no apparent superagonistic properties. We anticipate our tumor-selective agonist approach will have a more manageable safety profile, as it at least eliminates toxicity risks based on the engagement of TAA’s with low expression levels outside the tumor. Nonetheless, as with any drug in the CD28 agonist space, carefully designed clinical trials with low initial anti-PD-1 doses and protocols for the early detection and rapid management of immune-related adverse events are crucial.

Our CD28xVISTA bsAb was designed to complement PD-1/PD-L1 inhibitors, tumor-targeted radiolabeled therapies, or to enhance bispecific T-cell engagers’ selectivity and efficacy by targeting dual/orthogonal antigens on tumor and myeloid cells. A CD28xVISTA bsAb could be active in a broad set of solid tumors (as opposed to e.g. PSMA-targeting specifically in prostate cancer) and could be combined with any CD3xTAA TCE (or CAR-T cell therapy) in any solid tumor type with myeloid cell infiltration to potentially enhance their therapeutic window. Additionally, having both TCE engagement with a tumor antigen and CD28xVISTA bsAb engagement with VISTA on myeloid cells in the TME may be considered an “AND” logic gate for enhanced safety profile in combination treatment settings. With activity based on two components: TCE engagement with a TAA and CD28xVISTA bsAb engagement with VISTA at low pH and the functional assembly on the T-cell surface, this could generate a larger therapeutic window compared to TCE monotherapy.

Our results support the clinical testing of CD28xVISTA^pH-sens^ BS2 against cancers with tumor cells expressing VISTA as well as a broad variety of cancers with high level VISTA^+^ myeloid cell infiltration. We are currently exploring the use of pH-selective engagement of VISTA in the TME as a general platform for generation of tumor-specific bispecific antibody agonists for cancer therapy.

## METHODS

### Antibodies

TGN1412 was generated in-house from the published patent (55). CD28xVISTA bsAbs were based on a single CD28-binding Fab arm or disulfide-stabilized scFv derived from mAb TGN1412 (including a mutation changing a Cys sequence liability in VH CDR2 to Ser) and VISTA-binding Fab arms from anti-VISTA mAb 55873 (non-pH selective parental clone) or 67375 (pH-selective lead optimized version of 55873). Constructs with pH-selective VISTA binding domains from clone 67375 are indicated by “VISTA^pH-sens^” throughout the manuscript. These mAbs were obtained through selection of yeast-based platform libraries alternating between positive enrichment rounds at pH 6.0 and negative selection rounds at pH 7.4 (9). To ensure proper heavy and light chain pairing and purification of correctly formed homodimer we utilized the CrossMab (CH1-CL) technology (56) combined with Knob-side heavy chain (HC) mutations: T366W; disulfide stabilizing mutation: S354C; and Hole-side HC mutations: T366S/L368A/Y407V, disulfide stabilizing mutation: Y349C, and mutations: H435R/Y436F minimizing Protein A binding. Knob-in-hole mutations are referenced in (57); S-S-stabilization by (58,59), and mutations minimizing Protein A binding in (60). The CD28-binding scFv domain derived from TGN1412 was engineered for enhanced stability by introducing a disulfide bond between VH44 and VL100 (Kabat numbering; (61)). CD28xVISTA bsAbs all contain the L234S/L235T/G236R mutations silencing FcγR interactions (10). The comparator CD28^ADI^xVISTA^pH-sens^ BS2 maintains the same geometry as CD28xVISTA^pH-sens^ BS2 but differs in the anti-CD28 Fab arm (from an internal Adimab, LLC discovery program) along with proprietary Fc silencing and chain-pairing mutations. The CD28xPSMA comparator R-5678 was generated based on the variable regions of REGN5678 (employing a common LC for the CD28 and PSMA binding arm; (16) construct bs16429D) fused to IgG1 HC and Kappa LC with the HC pairing and Fc-null mutations mentioned above, and a common LC for the CD28 and PSMA binding arm. Synthetic genes carrying an N-terminal IL-2 signal peptide (GeneArt, Thermo Fisher Scientific) were cloned into the expression vector pcDNA™3.4 TOPO® (Thermo Fisher Scientific). Heavy and light chain sequences were co-expressed using the ExpiCHO Expression System (Thermo Fisher Scientific A29133) and antibody purification performed on a Protein A affinity column (HiTrap MabSelect SuRe^TM^ pcc resin; Cytiva 17549112 (62)) with 0.1M Glycine, pH 3.2 elution. After neutralization and pooling of main peak fractions, material was buffer exchanged into PBS (Lonza 17-517Q) using a μPulse Tangential Flow System (Formulatrix). An IEX polishing step was conducted as needed, and protein concentration of purified material was determined from the absorbance at 280 nm, where the molar extinction coefficient at 280 nm was calculated based on the amino acid sequence. Endotoxin levels were measured on an Endosafe nexgen-PTS instrument (Charles River). *In vivo* studies adhered to FDA guidance, maintaining endotoxin levels below 5EU/kg/h. For *ex vivo* CRS studies, endotoxin concentrations were kept below 0.01 EU/ml in blood, following Immuneed’s specifications. QC included analytical SEC with a typical monomeric purity of >95%. Specifically, for CD28xVISTA^pH-sens^ BS2, a SEC-HPLC purity of >99% monomer consisting exclusively of correctly paired heterodimer as analyzed by intact LC/MS was achieved after Protein A capture and a single IEX polishing step.

### Cell lines

Details about the source of the various cell lines are as follow: MC38 (Kerafast ENH204-FP), Jurkat (Clone E6-1; ATCC TIB-152), LNCaP (Clone FCG; ATCC CRL-1740) HEK-293 (ATCC CRL-1573), CHO-K1 (ATCC CCL-61), Kasumi-3 (ATCC CRL-2725). The Jurkat-IL-2-luciferase reporter cells were sourced from Promega (CD28 Bioassay, Core Kit; Promega JA6701). The CHO-K1/VISTA stable cell line was purchased from GenScript (GenScript M00533) HEK-293 cells were engineered by using the plasmid pcDNA3.1(+) (Thermo) for expression of membrane bound scFv derived from anti-CD3 clone OKT3 fused to the CD8 α-chain hinge and transmembrane domain or a fusion of this construct to the human VISTA gene through a furin cleavage site and T2A sequence allowing for co-expression of OKT3 scFv and VISTA from the same plasmid (63). The expression constructs were introduced into HEK-293 cells by Nucleofection (Lonza), and subsequent selection performed in Neomycin-containing media. MC38 cells overexpressing human VISTA were generated by transducing MC38 cells with lentivirus encoding a hVISTA-IRES-Puromycin gene fusion construct from an EF1α promoter (custom gene expression lentivirus; VectorBuilder). All other genes were codon optimized and custom synthesized by GeneArt (Thermo Fisher Scientific).

### Analysis of target binding

Recombinant protein binding to various antibodies was tested by ELISA experiments. 96-well flat bottom plates (Corning 2592) were coated with recombinant human VISTA-His (Sensei Biotherapeutics) or CD28-His (Acro Biosystems CD8-H52H3), 5 µg/ml in PBS pH 7.4 overnight at 4°C. The plates were blocked with PBS pH 7.4 or PBS pH 6.0 containing 2% NFDM for 2h at RT. The CD28xVISTA bsAbs as well as anti-VISTA and anti-CD28 monospecific control mAbs were three-fold serially diluted in PBS pH 7.4 or PBS pH 6.0 containing 1% NFDM starting at 300 nM before being added to the coated plates. Plates were incubated at RT for 2h. After washing 5x with PBS pH 7.4 or PBS pH 6.0 containing 0.05% Tween 20 (PBS-T), the antigen-antibody complexes were detected with a 1:60,000 dilution of goat-anti-human IgG-HRP (1h at RT) (Figure 1B & C). To show simultaneous binding to both targets (Figure 1D-F), antigen-antibody complexes were incubated with 100 ng/ml of biotinylated VISTA-His (Acro Biosystems B75-H82E1) in PBS pH 7.4 or PBS pH 6.0 containing 1% NFDM for 1h at RT followed by incubation with 1:400 dilution of Streptavidin-HRP (Pierce) in PBS pH 7.4 or PBS pH 6.0 containing 1% NFDM. Plates were washed 5x with PBS-T pH7.4 or PBS-T pH 6.0, and remaining HRP activity detected with TMB substrate (SeraCare). Optical density was measured at 450 nm. Data was analyzed by GraphPad Prizm version 10 using non-linear curve fitting (log[agonist] vs. response; variable slope (four parameters)). Binding of serial diluted CF647-labelled antibodies (Mix-n-Stain™ CF® Dye Antibody Labeling Kit, Biotium Cat. #92238) to native proteins on Jurkat T-cells (CD28), Kasumi-3 (VISTA) or a CHO cell line overexpressing VISTA were analyzed by flow cytometry. Briefly, 2.5×10^5^ cells were added to diluted antibodies in FACS buffer (MACSQuant® buffer, Miltenyi Biotec, Cat# 130-092-747) containing FcR Blocking Reagent (Miltenyi Biotec Cat. No. 130-059-901) and incubated for 20 minutes at 4°C. Cells were washed twice with FACS buffer and resuspended in 150 μL of FACS buffer containing 1μM SYTOX™ Blue dead cell stain (Thermo, Cat# S34857) for live/dead cell discrimination. Samples were analyzed using a MACSQuant Analyzer 10 flow cytometer (Miltenyi Biotec). Raw data were extracted using FlowJo and plotted using GraphPad Prism version 10. The number of surface-expressed VISTA molecules/cell for Kasumi-3 and CHO-K1/VISTA was determined using the Quantum™ Simply Cellular® (QSC) microsphere kit (Bangs Laboratories, Inc. 816) according to the manufacturer’s instructions.

### Reporter assays

BsAbs were tested for induction of luciferase expression from Jurkat-IL-2-luciferase reporter cells (CD28 Bioassay, Core Kit; Promega JA6701) in the presence of HEK-293 cells expressing membrane bound OKT3-scFv (anti-CD3) or OKT3-scFv+VISTA and CHO-K1 cells or CHO-K1 overexpressing human VISTA. The relevant HEK-293 and CHO-K1 cells were seeded at 20,000 cells each line/well and co-cultured in F12K medium with 10% FBS at 37°C, 5% CO_2_ overnight. After removing culture supernatant, cells were treated with CD28xVISTA BS1 in 40 µl/well media (RPMI-1640 with 10% FBS) at 0, 0.0003, 0.001, 0.003, 0.01, 0.04, 0.11, 0.33, 1, and 3 µg/ml for 1 hour at room temperature. A 40 μl/well suspension of TCR/CD3 Effector Cells (IL-2) was added at 80,000 cells/well, and cultured at 37°C, 5% CO_2_ for 5 hours. To determine the luminescence intensity, 80 μl/well Bio-Glo™ Reagent (Promega) was added to the plates and these were immediately read in a Molecular Devices Spectramax iD5 plate reader. The net relative luminescent unit (RLU) for each Ab-treated group was obtained after deducting the signal of non-Ab-treated controls.

### Assessment of CD28xVISTA^pH-sens^ BS2 in an *in vitro* Cytokine Release Assay

Cytokine release from human peripheral blood mononuclear cells (PBMCs), co-cultured with human umbilical vein endothelial cells (HUVECs) and treated with CD28xVISTA^pH-sens^ BS2, CD28^ADI^xVISTA^pH-sens^ BS2 (both with pH selective VISTA binding), R-5678 or TGN1412 as a positive control was examined. HUVEC:PBMC co-culture assays were conducted essentially as described in (19) using soluble antibody. HUVECs, allogenic to human PBMCs (Lonza C2519A), were expanded in Full EBM-2 Medium containing all BulletKit supplements (Lonza CC-3162). Cells were seeded into clear flat-bottom TC-treated 96-well plates (Fisher Scientific 3585) at a density of 30,000 cells/well in 100 μl medium. After culturing for 24h, the medium was replaced with human PBMC’s in full RPMI 1640 medium (Gibco A10491-01) containing 2% AB serum (Sigma H6914-100ML) and 1x non-essential amino acids (NEAA; Gibco 11140-050) (200 μl/well of 500,000 cells/ml). Antibodies were diluted and titrated to reach final assay concentrations of 1, 10, 30 and 100 µg/ml in full RPMI 1640, added to the plates (100 μl/well), and the co-culture incubated at 48h prior to cytokine analysis on a Bio-Rad Bio-Plex 200 instrument using the Bio-Plex Pro Human Cytokine 8-Plex Kit (Bio-Rad M50000007A) following the kit manual, with freshly prepared, reconstituted standards and positive controls diluted in Full RPMI-1640.

### Cytokine release profile of CD28xVISTA^pH-sens^ BS2 in an *ex vivo* system that mimics human blood circulation

*Ex vivo* cytokine release in human whole blood from 6 healthy donors by CD28xVISTA^pH-sens^ BS2, CD28^ADI^xVISTA^pH-sens^ BS2 (both with pH selective VISTA binding), and R-5678 was tested by Immuneed, AB using their ID.Flow circulating blood platform (20) with the controls anti-CD28 (ANC.28.1; 1 μg/ml), Alemtuzumab (3 μg/ml) or Cetuximab (250 μg/ml). Fresh whole blood was taken from healthy volunteers and a low amount of soluble heparin (allowing for analysis of drug-related effects on complement or coagulation cascade systems) was added. Blood was immediately transferred to the ID.Flow system, followed by administration of the test items, and set to circulate at 37°C to prevent clotting. Blood was extracted at baseline and at 4 hours and automatically counted using a Sysmex XN-L350 Hematology Analyzer. Cytokines in blood samples processed to plasma by centrifugation were measured using the Multi-Array platform from Meso Scale Discovery (MSD).

Proportions of T-cells (CD3^+^) and NK cells (CD56^+^ CD3^-^) expressing the activation marker CD69 was determined by flow cytometry using CD3-BV510 (BioLegend 300448), CD56-APC (Biolegend; 362504), CD69-PE/Cy7 (BioLegend 310912) and viability dye staining (LIVE/DEAD violet Viability Dye; Invitrogen L34964). Flow cytometry analysis was performed on a Cytoflex instrument (Beckman Coulter), and data analyzed using FlowJo v10.9.0.

### Effect of CD28xVISTA BS2 in LNCaP cell killing by CD3xPSMA TCE and primary human T-cells

Human T-cell mediated killing of LNCaP prostate cancer cells was analyzed on the xCELLigence real-time cell analysis platform (Agilent) by co-culturing LNCaP cells with PBMCs and VISTA^+^ Kasumi-3 cells in the presence of a CD3xPSMA bispecific T-cell engager (BPS Bioscience; Cat. # 101242-2 with a CD3-binding domain from the CD3xCD20 TCE Mosunetuzumab) alone or in combination with CD28xVISTA BS2. LNCaP cells were seeded at 10,000 cells/well into an E-plate (Agilent) and cultured in RPMI-1640 medium with 10% FBS for 3 days at 37°C, 5% CO_2_. After removing culture supernatant, medium with CD3xPSMA bsAb was added for a final concentration of 0, 0.001, 0.004, 0.01, 0.04, 0.11, 0.33, or 1 μg/ml. Human PBMCs (10,000 cells/well), Kazumi-3 cells (10,000 cells/well) and CD28xVISTA BS2 (0.1 μg/ml or medium in controls) was added to a final volume of 200 μl/well. Lysis buffer was added into control wells. The cells were cultured at 37°C, 5% CO_2_ and cell viability dynamically monitored on the xCELLigence instrument for 6 days. Only growth of the adherent LNCaP cells caused a change in the impedance signal measured by the instrument.

Cytokines in culture supernatant samples retrieved from the plate after 1 day were analyzed with a Bio-Plex Pro Human Cytokine 8-plex Kit (Bio-Rad). T-cell activation and proliferation were measured in samples retrieved after 6 days using flow cytometry on a Miltenyi MACSQuant Analyzer 10 instrument with CD4-VioBlue, CD8-VioGreen, CD25-PE, CD3-APC, and human Fc blocking reagent in MACS wash buffer with Propidium Iodide solution added immediately before sample analysis for live/dead cell discrimination (all flow reagents from Miltenyi Biotec).

### In vivo studies

Tumor growth inhibition (TGI) of a MC38 cell population overexpressing human VISTA was tested in a humanized CD28 mouse model (huCD28-Tg mice (C57BL/6N-*Cd28*^tm1.1(CD28)Geno^, genOway) in combination with anti-murine PD-1 (anti-mPD-1).

MC38-hVISTA cells were pre-conditioned by a passage through hCD28-KI mice followed by tumor harvesting and cell line establishment. Expression of hVISTA on preconditioned cells was analyzed by flow cytometry using fluorescently labelled anti-human VISTA mAb h26A (Sensei Biotherapeutics) prior to implantation. These MC38-hVISTA cells (1×10^6^/animal) were implanted subcutaneously into female hCD28 KI mice. Once the tumor volumes reached ∼80-100 mm^3^ (∼day 7) mice were randomized into 4 groups of 10 animals per group. Animals were administered isotype controls (rat IgG2a isotype control (clone 2A3, BioXCell BE0089), human IgG1 isotype control (BioXCell BP0297), anti-mPD-1 (InVivoMAb rat anti-mouse PD-1 (clone RMP1-14, BioXCell BE0146), CD28xVISTA^pH-sens^ BS2, or anti-mPD-1 + CD28xVISTA^pH-sens^ BS2 intraperitoneally twice/week for 3 weeks. Tumor growth inhibition was calculated as previously described (13). Significance was evaluated using Mann-Whitney unpaired t test with P < 0.05 considered to be statistically significant.

The PK-study was performed in a mixed population of female and male hCD28 KI mice (C57BL/6N-*Cd28*^tm1.1(CD28)Geno^; GenOway); n=4 per timepoint subgroup. The 3 CD28 targeting bsAbs CD28xVISTA^pH-sens^ BS2, CD28^ADI^xVISTA^pH-sens^ BS2 and R-5678 were dosed by bolus IV injection at 5 mg/kg. Blood was collected at 14 timepoints: pre-bleed; 5 min, 1, 4, 8, 12, 24, 48, 72, 120, 168, 336, and 672 hr, immediately put on ice and processed to serum which was frozen at −80°C until analysis. mAb serum levels were measured by ELISA using mouse anti-human IgG Fc (Abcam ab99757) immobilized in high bind microplates (Corning 2592) followed by detection using a peroxidase-conjugated mouse anti-human IgG F(ab)2 fragment-specific reagent (Jackson Laboratories 209-035-097; 40,000-fold diluted in Blocking Buffer PBS + 2% BSA) and development using the HRP substrate TMB (Life Technologies 34028). The concentrations in the samples were determined using non-linear regression with interpolation of unknown values from the prepared standard curve of the identical mAb using GraphPad Prizm 10 (GraphPad Software). Calculation of PK parameters using non-compartmental analysis was performed with Phoenix WinNonlin (version 8.3, Certara Corp.).

