## supplementary_info for "Dual Targeting for Enhanced Tumor Immunity: Conditionally Active CD28xVISTA Bispecific Antibodies Promote Myeloid-Driven T-Cell Activation"

Supplementary Information:

Supplementary Figures 1 – 2

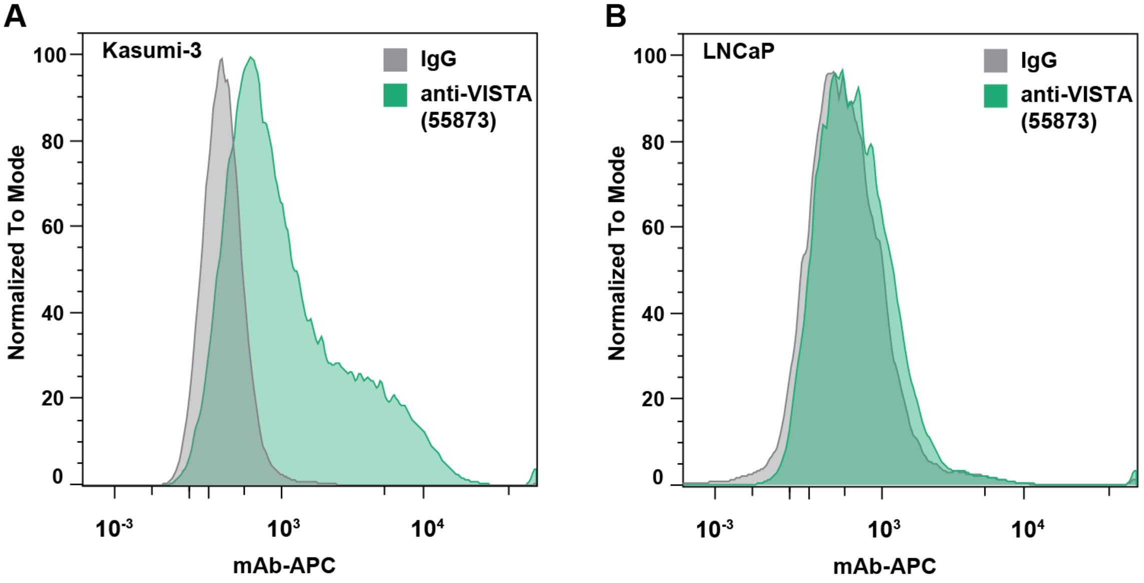


**Supplementary Figure 1: Expression of VISTA on Kasumi-3 and LNCaP cells. A,** Distribution of hVISTA+ Kasumi-3 human myeloid cells as measured by flow cytometry with a fluorescently labeled anti-hVISTA mAb (green) versus isotype control staining (gray). **B,** Distribution of hVISTA+ LNCaP human prostate cancer cells (green) versus isotype control staining (gray).

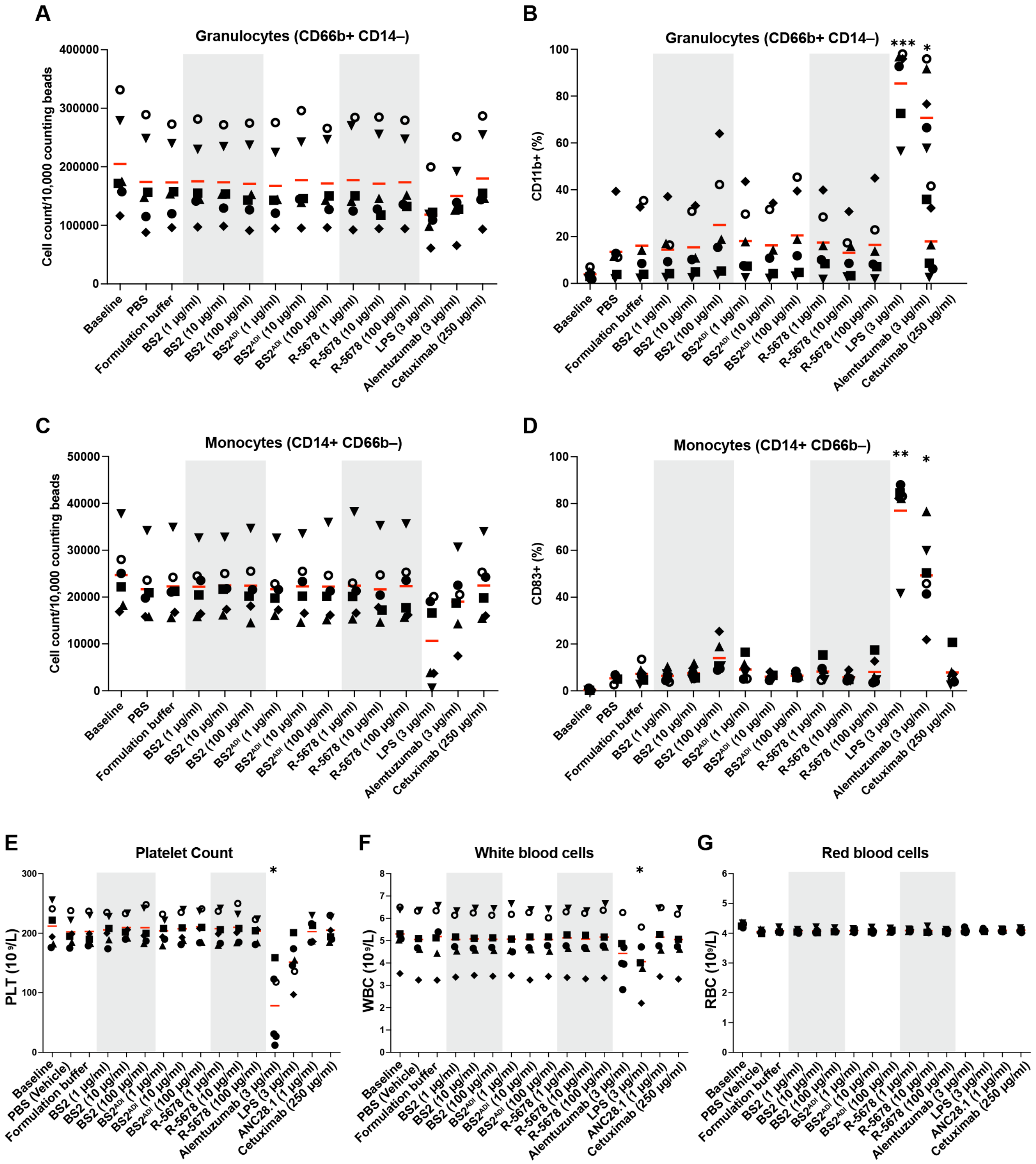


**Supplementary Figure 2: Full blood cell characterization from *ex vivo* human whole blood ID.Flow assay. A,** Granulocyte counts and **B,** activation (CD11b+) were not significantly affected by any of the test items at all three concentrations. **C,** Monocyte counts and **D,** activation (CD83+) were not significantly affected by any of the test items at all three concentrations. **E-G,** Indicated treatments had no effect on blood cell viability including **E**, platelet (PLT); **F**, white blood cell (WBC); and **G**, red blood cell (RBC) counts.

A single n= 1 experiment was performed with n= 6 independent donor samples. Symbols represent individual donor samples, mean values are indicated by red line. *p<0.05; **p<0.01; ***p<0.001; ****p<0.0001 comparison to formulation buffer (**B** through **G**) by Paired Student's t-test with Holm-Sidak correction. BS2: CD28xVISTA$^{pH\text{-}sens}$; BS2$^{ADI}$: CD28$^{ADI}$xVISTA$^{pH\text{-}sens}$; R-5678: CD28xPSMA.
